# Mortality causes and annual survival rates of red foxes *Vulpes vulpes*

**DOI:** 10.64898/2026.07.18.739295

**Authors:** Tomas Willebrand, Morten Odden, Zea Walton, Gustaf Samelius, Kjartat Østbye, Bo Soderberg, Goran Spong

## Abstract

Mortality causes and survival rate estimates from individual follow-ups of GPS-collared animals provide reliable demographic data, but such estimates are rare for red foxes *Vulpes vulpes*. We used data from 126 GPS-collared red foxes tracked between 2011 and 2019 across a latitudinal gradient in central Sweden and Norway to quantify mortality causes and estimate annual survival probabilities using the Andersen-Gill extension of the Cox proportional hazards model. Hunting was the dominant mortality cause (63%), followed by vehicle collisions (14%), stress or malnutrition (11%), sarcoptic mange (9%), and predation (3%). Annual survival was 0.58 for adults and 0.32 for subadults. Subadults had approximately twice the hazard of adults, males had elevated hazard relative to females, and mortality risk was highest in autumn and early winter. These collar-based estimates are strikingly similar to mark-recovery estimates from the same region nearly five decades earlier, despite an intervening population collapse from sarcoptic mange, subsequent full recovery, and a major decline in harvest pressure. This convergence is consistent with density-dependent regulation at a food-determined and food-limited carrying capacity, suggesting that current harvest levels are not limiting the population. Furthermore, human-caused mortality was approximately four times greater than natural mortality, illustrating that living in close proximity to humans — which often favours generalist predators such as red foxes — may also come at a cost of increased mortality risk. Developing validated census methods for red foxes in boreal Scandinavia is identified as a key priority for quantitative population management.

## Introduction

Management models based on demographic mechanisms have the potential to identify the relative importance of survival, productivity and recruitment for population change, and therefore better support management decisions (Lebreton and Gimenez, 2013; Gaillard et al. 2003; Sæther and Bakke, 2000). Unfortunately, obtaining data on survival and reproductive traits are difficult for most wildlife, and estimates of demographic rates often suffer from small samples that results in large uncertainties (Doak et al. 2005; Popescu et al. 2016). Survival is particularly difficult to obtain, and was traditionally estimated from mark-recapture analyses and life-table analysis based on age-at-harvest. However, continuous tracking of radio-collared individuals provides a reliable alternative to estimate survival time and mortality causes (White and Garrott, 2012), and survival estimates based on individual follow-ups are more reliable compared to other methods (Lebreton, Pradel, et al. 1993). These data can be used to estimate survival probabilities through Kaplan-Meier estimates (Pollock et al. 1989), and evaluate the effects of covariates in regression models such as the Andersen-Gill model (Andersen and Gill, 1982; Murray and Patterson, 2006).

The red fox *Vulpes vulpes* is a generalist predator that tends to benefit from human activities and settlements (Panek and Bresiski, 2002; Jahren et al. 2020; Díaz-Ruiz et al. 2016). The red fox is also considered a nuisance species due to its potential to spread disease that can affect humans (Anderson et al. 1981; Maas et al. 2014), and its predation on domestic, economically valuable and threatened species (McLeod et al. 2010; Jarnemo and Liberg, 2005; Kämmerle and Storch, 2019). Because of this, red foxes are often the target of control programs at different scales in large parts of its natural distribution (Rushton et al. 2006; Kämmerle and Storch, 2019; Loonstra et al. 2024; Heydon and Reynolds, 2000). However, the effectiveness of control programs is debated, and there are several knowledge gaps needing to be filled to better understand the population dynamics of red foxes and the potential effects of control. There is a lack of reliable estimates of population density and survival probabilities from individual follow-ups are rare. These data are particularly important given their ability to rapidly recolonize areas after control efforts cease (Newsome et al. 2014).

Winter conditions at northern latitudes are limiting (Barto and Zalewski, 2007), but the red fox has expanded north as human settlements and infrastructure developed in the tundra and arctic regions (Gallant et al. 2020). In this study we aim to identify 1) the importance of different mortality causes, 2) estimate survival probabilities, and 3) identify the effects of age, sex and season on hazard rates. We use data from 126 GPS-collared red foxes in Sweden and Norway and expect males and subadults to have lower survival probabilities compared to females and adults. We expect subadults to be vulnerable due to inexperience and dispersal behavior, and males to be more vulnerable than females due to their larger home ranges and more exploratory behavior (Walton, Samelius, et al. 2017; Englund, 1980; Walton, Hagenlund, et al. 2021). Due to their association with human settlements and infrastructure, we expect exposure to anthropogenic mortality causes to be common, and that seasonal patterns in mortality causes and hazard rates increase in autumn/winter during the hunting season.

## Methods

### Study Area

Foxes were captured and collared in four different areas along a gradient of decreasing landscape productivity and human land use as latitude increased from 58^◦^ to 62^◦^ N in central Sweden and Norway. The southernmost area consists of fragmented boreonemoral forests, agricultural lands, and human settlements, whereas the northern areas are dominated by boreal forests and scattered human settlements. Norway spruce *Picea abies*, Scots pine *Pinus sylvestris*, and to a lesser degree birch *Betula* spp. are the most common tree species. Broad-leaved deciduous species such as English oak *Quercus robur* and Norway maple *Acer platanoides* are present in the southern area but rare in the north. The northernmost part grades into alpine tundra of low diversity and productivity. Altitude increases with latitude from approximately 25 to 750 m above sea level, and snow cover shifts from irregular in the south to continuous from November through May in the north. For further details see Walton, Samelius, et al. (2017).

### Capture, Tagging & Tracking

All captured foxes heavier than 5 kg were fitted with a GPS radio collar (Tellus Ultra-light, 210 g, Televilt Inc., Lindesberg, Sweden). Foxes were sexed, measured, weighed, and aged at capture. Age was classified as subadult (< 1 year) or adult (> 1 year) based on the degree of tooth wear and tooth coloration. GPS collars were programmed to acquire a minimum of three positions per day; variation in duty cycles resulted in collar drop-off after 6–9 months. All capture and handling procedures were approved by the Swedish Animal Ethics Committee (permit numbers DNR 70–12, DNR 58–15) and the Norwegian Experimental Animal Ethics Committee (permit numbers 2009/122825, 2012/20038, 2014/207803). Permits to capture wild animals were issued by the Norwegian Directorate for Nature Management and the Swedish Environmental Protection Board (NV-03459-11). For further details see Walton, Samelius, et al. (2017) and Walton, Samelius, et al. (2018).

Between 2011 and 2019, we captured and GPS-collared 126 individual red foxes. Six individuals were recaptured after a monitoring gap, yielding 132 capture events in total. One fox (Ann) was tracked continuously using two successive collars and its records were merged into a single uninterrupted interval. Five foxes (Frans, Gunilla, Ingvar, Oskar, Cristina) were recaptured after a gap in monitoring and treated as two independent observation intervals, consistent with the Andersen-Gill counting-process framework (Andersen and Gill, 1982). Cristina was subadult at first capture and adult at recapture, and her two intervals were assigned the appropriate age class separately. We excluded four intervals (Gunilla’s first capture, Källsvedan, Oskar’s second capture, and Stiga) that contributed fewer than two weeks of tracking and were therefore uninformative for survival estimation, leaving a total of 127 individual fox-intervals in the analysis (Table 1).

**Table 1:** Number of collared fox intervals included in the analysis, by age class and sex. One fox (Ann) contributed as a single merged interval across two collars; five foxes contributed two intervals each following recapture. Four individuals tracked for fewer than two weeks were excluded.

| Sex | Adult | Sub-adult | Total |
| --- | --- | --- | --- |
| Females | 28 | 22 | 50 |
| Males | 44 | 33 | 77 |
| Total | 72 | 55 | 127 |

**Table 2:**
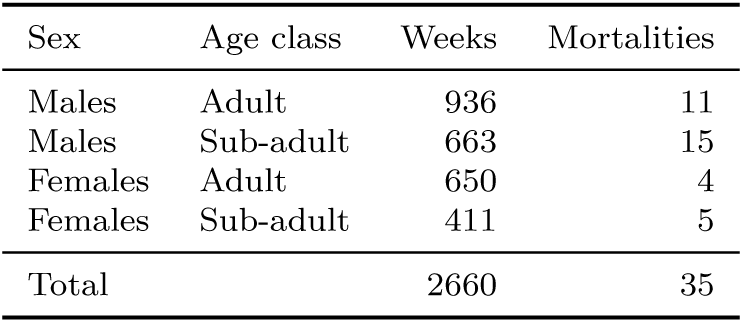
A summary of counting-process dataset in which each row represents the interval (*t*_start_*, t*_end_] for that individual-week. The last week a fox was monitored was recorded as either a known mortality or a censored observation. In total, the 127 fox individuals provided 2660 fox weeks and 35 mortality events. The table is stratified by sex and age class (adult vs. sub-adult), and the last column shows the number of mortality events in each stratum.

| Sex | Age class | Weeks | Mortalities |
| --- | --- | --- | --- |
| Males | Adult | 936 | 11 |
| Males | Sub-adult | 663 | 15 |
| Females | Adult | 650 | 4 |
| Females | Sub-adult | 411 | 5 |
| Total |  | 2660 | 35 |

An inactive GPS collar transmitted a mortality signal and SMS notification used to record the approximate time of death. Mortality causes other than hunting were determined in the field: damage consistent with vehicle strikes was assessed visually; carcasses exhibiting alopecic patches, crusted skin, and sometimes a foul aromatic odour were classified as sarcoptic mange; and a wolf kill was identified by tracks and bite marks at the kill site. Most mortalities resulted from hunting, and hunters returned collars with the date of harvest. Cases where cause of death could not be determined were classified as unknown.

### Analysis

To estimate the effects of age, sex, and season on survival we used the Andersen-Gill extension of the Cox proportional hazards (CPH) regression model (Andersen and Gill, 1982), which can handle time-varying covariates and recaptures (Johnson et al. 2004; Åhlen et al. 2013). The non-parametric Kaplan-Meier estimator was used to visualize and inspect survival curves (Pollock et al. 1989).

Each fox contributed one row per calendar week it was present in the study, yielding a counting-process dataset in which each row represents the interval (*t*_start_, *t*_end_] for that individual-week. The last week a fox was monitored was recorded as either a known mortality or a censored observation due to collar drop-off or failure; all preceding weeks were censored. Collar failure and collar drop-off (CAUSE = 9) were treated as non-informative right-censoring throughout. We pooled study areas and years due to low overall sample size.

We set the start of the red fox year to ISO week 36 (early September), when cubs typically separate from family groups, and expressed time within the year as a continuous fox-week variable running from 1 (week 36) to 52 (week 35 of the following calendar year). Sub-adults whose tracking extended beyond week 35 of the calendar year following their capture year were reclassified as adults from that point onward; two individuals (Cristina and Gijom) met this criterion.

Robust sandwich standard errors, obtained via cluster(FoxID) in the coxph func-tion, accounted for the non-independence of multiple observation rows from the same individual (Therneau, 2024). We developed and compared five candidate CPH models containing combinations of age (subadult vs. adult), sex, and season (Table 4). Season was evaluated as: (i) a binary variable (autumn/winter vs. spring/summer), (ii) a continuous linear fox-week covariate, (iii) a sine-cosine pair capturing circular annual periodicity, and (iv) a four-level factor (autumn, winter, spring, summer). Models were compared using Akaike’s Information Criterion (AIC) computed from the partial log-likelihood. Likelihood ratio tests (LRT) between nested models were performed on refitted models without the cluster() argument, because the robust variance estimator does not alter the partial log-likelihood and therefore does not affect LRT statistics. We assessed whether a sex × season interaction improved model fit, but given only nine female mortality events and the rule of thumb of approximately ten events per estimated parameter, this interaction was considered underpowered and is reported for transparency only.

The proportional hazards assumption was evaluated using scaled Schoenfeld residuals via cox.zph with a rank time transformation (Therneau, 2024). All data preparation and analyses were conducted in R version 4.6 (R Core Team, 2024), using the packages survival (Therneau, 2024) and survminer (Kassambara et al. 2024).

## Results

### Mortalities

We registered 35 known mortality events from the 126 collared foxes. *Harvest* was the most common mortality cause (63%, n=22), followed by *vehicle collision* (14%, n=5). Three (9%) died as consequences of *sarcoptic mange*, one (3%) was killed by *wolf predation* but left intact. The four foxes (11%) categorized as *stress/malnutrition* were uncertain causes and contain suspected stress, malnutrition or disease. Harvest did not appear to be a more common source of mortality in males compared to females, 62% to 67% (Fisher’s exact test, P = 1, Odds ratio = 0.92). However, four of the five recorded mortalities of subadult females were attributed to hunting. See table 3. We note that all vehicle deaths and all stress/malnutrition deaths were male, and all disease deaths were female, although this should be interpreted cautiously given the limited sample size.

**Table 3:** Mortality causes of 35 recorded deaths from 126 collared foxes separated by age and sex.

| Age | Sex | Hunting | Traffic | Disease | Wolf kill | Stress / Malnutrition | Total |
| --- | --- | --- | --- | --- | --- | --- | --- |
| Adult | Female | 2 | 0 | 2 | 0 | 0 | 4 |
| Adult | Male | 8 | 2 | 0 | 0 | 1 | 11 |
| Sub-adult | Female | 4 | 0 | 1 | 0 | 0 | 5 |
| Sub-adult | Male | 8 | 3 | 0 | 1 | 3 | 15 |
| Total |  | 22 | 5 | 3 | 1 | 4 | 35 |

### Survival

Kaplan-Meier curves revealed that sub-adults had markedly lower survival in early autumn compared to adults, with the curves converging over time (Figure 1). The log-rank test indicated a significant difference in survival between age classes (p=0.043) and a marginally non-significant difference between sexes (p=0.095). However, these curves do not adjust for covariates and should be interpreted as visual tools rather than definitive evidence of differences.

**Figure 1:**
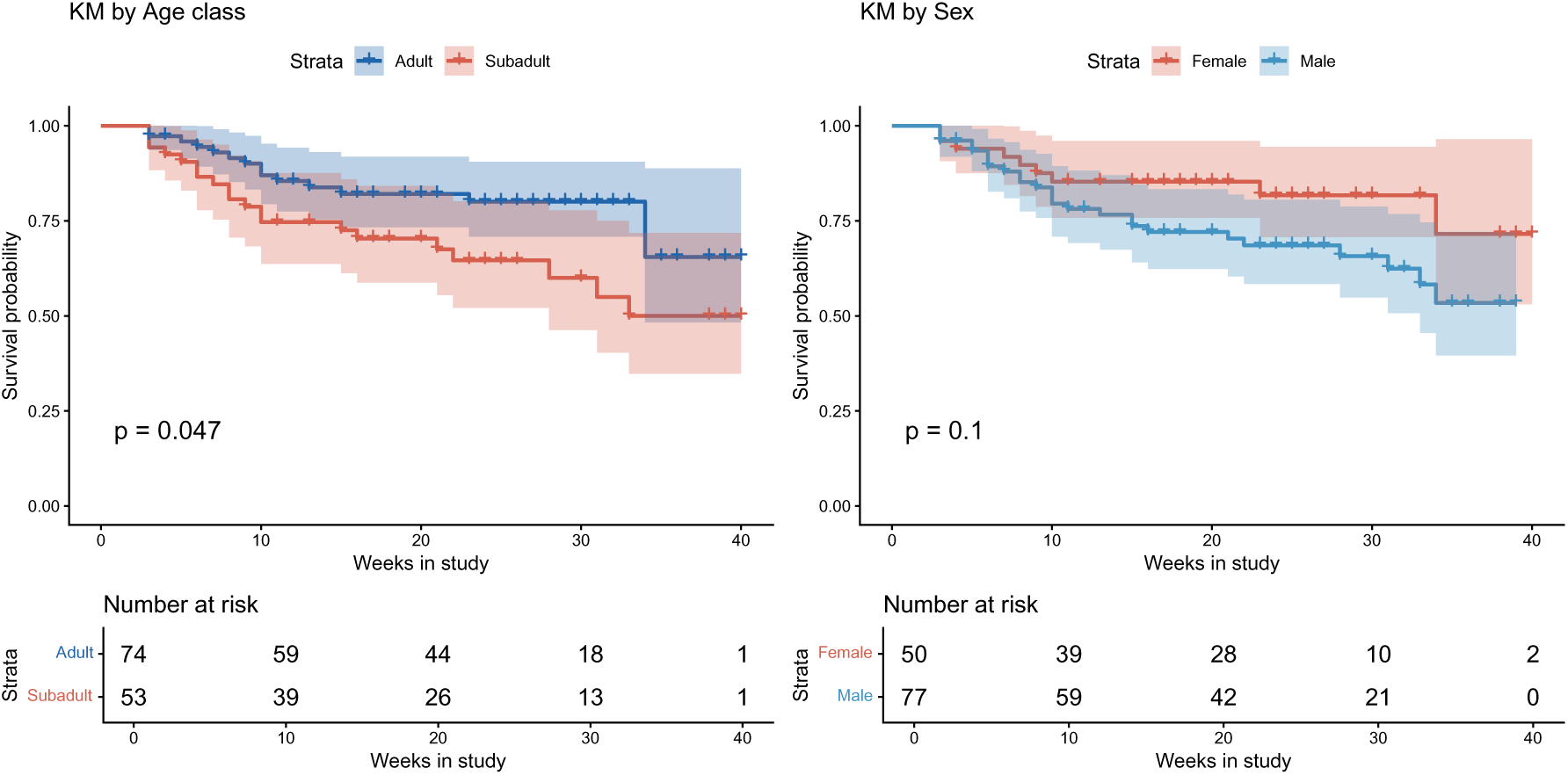
The figure shows Kaplan Meier survival curves stratified by age (left) and sex (right). The y-axis shows the survival probability and the x-axis shows time in fox-weeks, which starts at ISO week 36 (years pooled). P-values shown are the results of the log-rank test. The vertical lines on the survival curves indicate censored observations or mortality events. The tables below the graphs show the number of individuals at risk at each time point.

**Figure 2:**
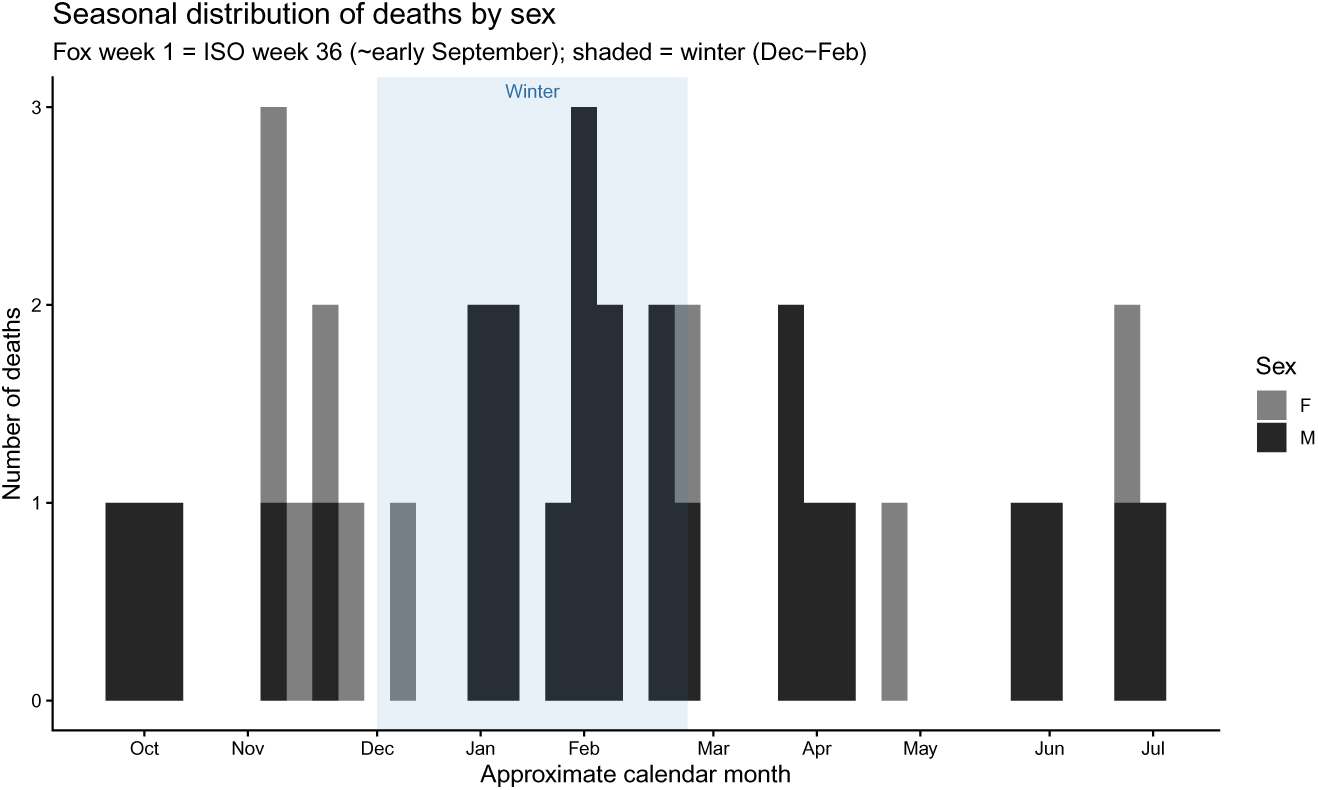
The figure shows the seasonal distribution of deaths stratified by sex, males=black and females=grey. The y-axis show the frequency of mortalities and the x-axis shows the month of the year. A higher frequency in autumn/winter (blue shade).

The five different Cox PH models are compared in Table 4. The model including age, sex, and season as a continuous linear fox-week covariate provided the best fit (lowest AIC, concordance = 0.66). The binary season coding performed worst, and adding season improved concordance substantially over the age+sex base model. The sex × season interaction did not improve model fit (LRT *χ*^2^ = 1.10, df=1, p=0.295), consistent with insufficient power to detect a sex-specific seasonal pattern given only nine female mortality events.

**Table 4:** The table presents a comparison of the different Cox proportional hazards models evaluated. The first column lists the model specifications, the second column shows the Akaike Information Criterion (AIC) for each model, the third column shows the concordance statistic (C-index) which measures the predictive accuracy of the model, and the last column shows the number of parameters in each model. The best performing model based on AIC is highlighted in bold.

| Model | AIC | Concordance | Params |
| --- | --- | --- | --- |
| Age only | 307.26 | 0.587 | 1 |
| Age + Sex | 306.68 | 0.618 | 2 |
| **Age + Sex + Season (linear)** | **306.05** | **0.658** | **3** |
| Age + Sex + Season (binary) | 308.67 | 0.618 | 3 |
| Age + Sex + Season (4-factor) | 306.64 | 0.697 | 5 |

**Table 5:** The table compares the hazard ratios of the three Cox proportional hazards models. The first column lists the model specifications, the second column shows the covariates included in each model, and the remaining columns display the hazard ratios (HR), confidence intervals (CI), and p-values. Age and sex are binary variables, while season is a continuous variable. The prefered model is the model with age, sex and season as a linear term.

| Model | Covariate | HR | CI_lower | CI_upper | p |
| --- | --- | --- | --- | --- | --- |
| Age only | AGE | 1.939 | 1.002 | 3.753 | 0.049 |
| Age + Sex | AGE | 1.910 | 0.988 | 3.691 | 0.054 |
| Age + Sex | SEX | 1.816 | 0.848 | 3.888 | 0.125 |
| Age + Sex + Season (linear) | AGE | 1.835 | 0.951 | 3.541 | 0.070 |
| Age + Sex + Season (linear) | SEX | 1.760 | 0.821 | 3.770 | 0.146 |
| Age + Sex + Season (linear) | season | 0.976 | 0.948 | 1.005 | 0.109 |

The preferred Cox PH model revealed that subadults had approximately twice the hazard of adults (HR 1.9, CI 0.884.2), though the confidence interval overlapped 1, re-flecting low precision. Males had elevated hazard relative to females (HR 1.8, CI 0.734.6), similarly imprecise. Hazard declined as the fox year progressed (HR per fox-week 0.96), suggesting autumn/early winter is the most dangerous period. Wide confidence intervals reflect the limited number of events (n=35) and that conclusions about direction of effects are more reliable than conclusions about magnitude.

The proportional hazards assumption was satisfied for all covariates in the preferred model (Schoenfeld residuals, all p > 0.4), supporting the validity of the CPH framework.

## Discussion

Our GPS-collar survival estimates (adults: 0.58, sub-adults: 0.32) are remarkably similar to the pre-mange mark-recovery estimates from Willebrand et al. (2022) of 0.55 and 0.36, respectively, despite being separated by nearly five decades that included a population collapse, full recovery, and substantial changes in the socioeconomic context of fox hunting. The collapse of the fur trade since the 1970s has greatly reduced the economic incentive for intensive harvest, which was a major component of harvest mortality in the pre-mange period. Although we cannot directly quantify the change in per-capita harvest rate over time without concurrent population size estimates, the convergence of survival estimates across these two very different periods is consistent with density-dependent regulation of survival at a food-determined carrying capacity, rather than with harvest as a primary limiting factor. If harvest were limiting, reduced harvest pressure would be expected to produce detectably higher survival in the current population a pattern not observed here. Instead, the population appears to return to a similar demographic equilibrium regardless of harvest intensity, driven by food availability and the resulting intraspecific competition for territories and resources. This interpretation is consistent with previous work which concluded that fox density in boreal Sweden tracks vole abundance closely (Englund, 1970; Lindström, 1989).

Mortality causes and survival estimates based on individual fates of red foxes are almost non-existent (but see Gosselink et al. (2007)), and our study provides an important contribution to the very limited literature on red fox demography. Our results are consistent with earlier studies based on mark-recovery and life-table analyzes, which also found that hunting is the most common mortality cause, followed by vehicle collisions and disease (Gosselink et al. 2007; Heydon and Reynolds, 2000; Yoneda and Maekawa, 1982; Goszczyski, 1989; Harris and Smith, 1987; Willebrand et al. 2022). The red fox is a species that is closely associated with human settlements and infrastructure (Jahren et al. 2020), and it is therefore not surprising that anthropogenic mortality causes are common in fox populations. However, the fact that human-caused mortality in our study was approximately four times greater than natural mortality suggests that living close to humans comes at a cost of increased mortality risk. In this study, hunting dominated the mortality causes despite extensive road networks and a persistent prevalence of sarcoptic mange in the red fox population (Carricondo-Sanchez et al. 2017). Gosselink et al. (2007) identified sarcoptic mange as the most common mortality cause in urban areas, with a similar prevalence level as in our area, 3.15% (average) versus 1–7% (range). In rural areas on the other hand, coyote predation had become the most important source of mortality after an estimated 10-fold decrease in hunting mortality. In Sweden, hunting red foxes is considered a valuable sport hunting activity as well as a source of pelts, and between 50 000 and 80 000 foxes have been harvested annually since the turn of the century (*Viltdata* 2025). Furthermore, Sweden lacks a predator equivalent to the coyote in the meso-predator guild. Large predators in Sweden that potentially kill red foxes are wolves *Canis lupus*, lynx *Lynx lynx*, wolverines *Gulo gulo* and golden eagles *Aquila chrysaetos* (Helldin et al. 2006; Sulkava et al. 1999; van Dijk et al. 2008), but these predators occur at low densities, especially close to human settlements. The red fox shows pronounced vigilant behavior (Berger-Tal et al. 2009), and camera traps placed at slaughter remains have shown that red foxes avoid approaching the site until scavengers such as wolverine and golden eagle have reduced their use (Gomo et al. 2017). We note that all vehicle deaths and all stress/malnutrition deaths being male, and disease deaths being female, could be a spurious pattern due to the limited sample size, but if real, it could be linked to the higher activity and exploratory behavior of males, and the higher vulnerability of females to sarcoptic mange due to their higher survival and therefore longer exposure time.

However, robustly testing whether harvest is additive or compensatory to other sources of mortality, and whether productivity or survival is the primary driver of population dynamics requires reliable estimates of population density. Currently, no validated census method exists for red foxes in the boreal and sub-arctic landscapes of Scandinavia. Camera trapping, scat collection and mark-recovery methods, and occupancy modelling based on non-invasive sampling have been validated for other carnivores in similar landscapes. Developing and validating such a method that can be integrated with existing wildlife monitoring is one of the most important steps to base fox population management on quantitative demographics. Without it, survival estimates from telemetry studies and age-at-harvest models remain difficult to interpret in terms of absolute population change, and the relative roles of harvest, food availability, and disease in driving fox population dynamics will be difficult to disentangle.

Although our sample size was limited, the results suggest that survival of sub-adults was lower than adults, and males had lower survival than females. The higher vulnerability of subadults was likely due to inexperience and higher dispersal rate than adult foxes, which is consistent with earlier studies (Gosselink et al. 2007; Yoneda and Maekawa, 1982; Goszczyski, 1989; Harris and Smith, 1987; Willebrand et al. 2022). A concurrent life-table analysis of 6 022 harvested red foxes from the same study system also showed that females had consistently higher survival than males (Willebrand et al., unpublished). Higher survival of females compared to males was also reported by Heydon and Reynolds (2000), although at lower survival rates than in our study. Seasonal patterns in mortality causes and survival rates in our study suggest higher mortality in autumn/winter, potentially linked to hunting seasons and resource scarcity. Together with the declining survival documented during the first mange outbreak (Willebrand et al. 2022) and the latitudinal patterns in age- and weight-dependent survival (Willebrand et al., in prep), these estimates provide a multi-decadal and spatially replicated demographic picture of red fox populations in boreal Scandinavia.

Several studies of harvested foxes have found a male bias in the samples (Harris and Smith, 1987; Gortázar et al. 2003; Goszczyski, 1989; Yoneda and Maekawa, 1982; Englund, 1970; Hartová-Nentvichová et al. 2010). The hunting season overlaps with the period when males show higher activity and move longer distances compared to females (Englund, 1980). In addition, adult males have larger home ranges than adult females, and tend to make more frequent and longer excursions compared to females (Soulsbury et al. 2011). The bias is reduced in intensively hunted areas, and even reversed when foxes are killed in spring (Yoneda and Maekawa, 1982; Goszczyski, 1989; Galby and Hjeljord, 2010). Higher survival rates of females will result in an excess of adult females in the population, which can explain why some adult males show exploratory movements and non-overlapping clusters in spring. A female-biased sex ratio in the red fox population likely increases the competition for mates among males and could further promote a risk-prone behavior of males (Clutton-Brock and Parker, 1992; Weir et al. 2011).

### Assumptions and limitations

Survival analysis is dependent on number of mortality events, and our sample size of 35 events limits precision and power to detect effects, especially for sex-specific patterns. The Cox model assumes proportional hazards, which we verified, but unmeasured confounders could bias results. Cause of death classification may be imperfect, particularly for unknown cases. Our sample is pooled from a large geographical region and may not be representative of all fox populations, limiting generalizability. Area and yearly differences will reduce precision of the parameter estimates of the Andersen-Gill model evaluating differences in sex and age. The confidence intervals around the survival estimates are large, but are comparable to estimates based on life-table models (Lebreton, Pradel, et al. 1993). We believe our result is a useful contribution when developing both theoretical and applied population models of red fox dynamics (Devenish-Nelson et al. 2013; Schaub and Kéry, 2021). We used lightweight GPS-collars (210 g) weighing less than 4% of the minimum body weight of the collared foxes (>5 kg), which is within typical guidelines for collar-to-body weight ratios.

## Conclusions

Anthropogenic mortality causes, particularly hunting, were the dominant sources of mortality for red foxes in our study system, and mortality risk was elevated in autumn/winter linked to hunting seasons and resource scarcity. However, one cannot conclude that present harvest rates limit the population, and we believe the convergence of survival estimates across decades suggests a density-dependent regulation at a food-determined and food-limited carrying capacity. We propose that developing and validating a method for estimating population density is prioritized to understand the relative roles of factors driving fox population dynamics. Density estimates would allow for testing whether harvest is additive or compensatory to other sources of mortality, and provide important data when developing integrated population models to support management decisions.

## Acknowledgements

We are grateful to students and volunteers that provided invaluable help in the field. Funding for this study was provided by the Swedish Environmental Protection Agency, the Swedish Hunters Association, Karl Erik Önnesjös Stiftelse (Sweden), and the Norwegian Directorate for Nature Management and the Gotaas Fund (Norway). All animal capture and handling protocols were approved by the Swedish Environmental Protection Board and the Swedish Animal Ethics Committee (permit numbers NV-0345911, DNR 7012, DNR 5815, DNR 1347).

## Author contributions

Tomas Willebrand: Conceptualization (equal); Data curation (lead); Formal analysis (lead); Funding acquisition (equal); Investigation (equal); Methodology (lead); Writing original draft (lead); Writing review and editing (lead). Gustaf Samelius: Conceptualization (equal); Formal analysis (equal); Investigation (equal); Methodology (equal); Validation (equal); Writing review and editing (equal). Zea Walton:Project administration (lead); Conceptualization (equal); Formal analysis (equal); Investigation (equal); Methodology (equal); Validation (equal); Writing review and editing (equal). Morten Odden: Conceptualization (equal); Investigation (equal); Methodology (equal); Valida-tion (equal); Writing review and editing (equal). Kjartan Østbye: Validation (equal); Writing review and editing (equal). Göran Spong: Validation (equal); Writing review and editing (equal). Bo Söderberg: Investigation (equal); Methodology (equal).

## Data availability

Data on fates and time present in the study used in this analysis can be found at Zen-odo.org (doi:10.5281/zenodo.21424954).

